# High prevalence of haemosporidian parasites in Eurasian Jays

**DOI:** 10.1101/2023.06.20.545710

**Authors:** Yvonne R. Schumm, Naemi Lederer-Ponzer, Juan F. Masello, Petra Quillfeldt

## Abstract

Avian haemosporidians are vector-borne blood parasites, which infect a great variety of avian host species. The order Passeriformes has the highest average infection probability; nevertheless, some common species of this order have been rather poorly studied in terms of haemosporidian prevalence and diversity. In this study, we investigated the prevalence of haemosporidians in one of such species, the Eurasian Jay *Garrulus glandarius* (Corvidae), from a forest population in Hesse, Central Germany. All individuals were infected with at least one haemosporidian genus, i.e. an overall prevalence of 100%. The most common infection pattern was a mixed infection with *Haemoproteus* and *Leucocytozoon*, whereas no *Plasmodium* infection was detected. Our results regarding lineage diversity combined with data from other studies indicate a rather pronounced host-specificity of *Haemoproteus* and *Leucocytozoon* lineages infecting birds of the family Corvidae.

## Introduction

Vector-borne avian blood parasites (Apicomplexa, order Haemosporidia), including the genera *Plasmodium* (avian malaria), and malaria-like pathogens *Leucocytozoon* and *Haemoproteus*, are widespread and infect a great variety of avian host species (Valkiūnas 2005). The application of molecular techniques for detecting and characterizing haemosporidians are providing an increasingly growing and accurate picture of their host range, (host-specific) prevalence, distribution, and diversity (Rivero & Gandon 2018). The intensified research has revealed an extensive genetic diversity of haemosporidian lineages (> 4800 lineages, MalAvi 2023), which seems to be matched by an equally rich phenotypic diversity, such as lineages differing in their host range or within-host effects or virulence (Rivero & Gandon 2018; Ágh et al. 2022). At avian host-species level, differences in host life history and behavioural traits, e.g. preferred foraging habitat or nest type, can influence rates of dipteran vector exposure, which can lead to heterogeneous infection probabilities across avian hosts (Fecchio et al. 2021; Rodríguez-Hernández et al. 2021). In a global study (Fecchio et al. 2021), including 141 avian families, those with the highest average *Leucocytozoon* infection probabilities belonged the order Passeriformes, e.g. Paridae, Corvidae. Also, the highest *Plasmodium* lineage diversity across all continents was found in passerine birds (Rivero & Gandon 2018). Nevertheless, in species of the order Passeriformes, many lineages and lineage-host relations likely were not found yet, particularly in hosts that have not been intensively sampled. For instance, there are only 23 entries for Eurasian Jays *Garrulus glandarius* in the MalAvi database (274 entries in total for Corvidae), comprising seven *Haemoproteus*, two *Plasmodium* and five *Leucocytozoon* lineages (MalAvi 2023). The Palearctic-oriental distributed Eurasian Jay is in Central Europe a resident, mainly forest-dwelling, open-nesting species, which in some years show eruptive movements in the autumn, which seem to be related to population density and the variation in acorn availability (Glutz von Blotzheim & Bauer 1992; Selås 2017). In this study we aimed to (a) assess the Haemosporidia infection status in Jays from one forest population in Central Germany, (b) investigate the lineage diversity, and (c) compare our results with known lineages in Eurasian Jays from other breeding sites.

## Material & Methods

### Blood sample collection

We sampled a total of 16 Eurasian Jays from May 2020 to July 2021. We captured the individuals using walk-in traps placed in the Marburg Open Forest, a 250-ha managed forest, consisting of mixed stands dominated by Common Beech *Fagus sylvatica* and Common Oak *Quercus robur*, in Hesse, Central Germany (50°50′ N, 8°39′ E). We obtained the blood samples by puncture of the brachial vein (Table 1), and stored them on Whatman FTA cards (Whatman®, UK). Additionally, we prepared two blood smears per individual, which we fixed with methanol (100%) for 30 s and stained with Giemsa in a work solution prepared with buffer pH 7.0 (ratio 1:5) for 30 min.

**Table 1.**
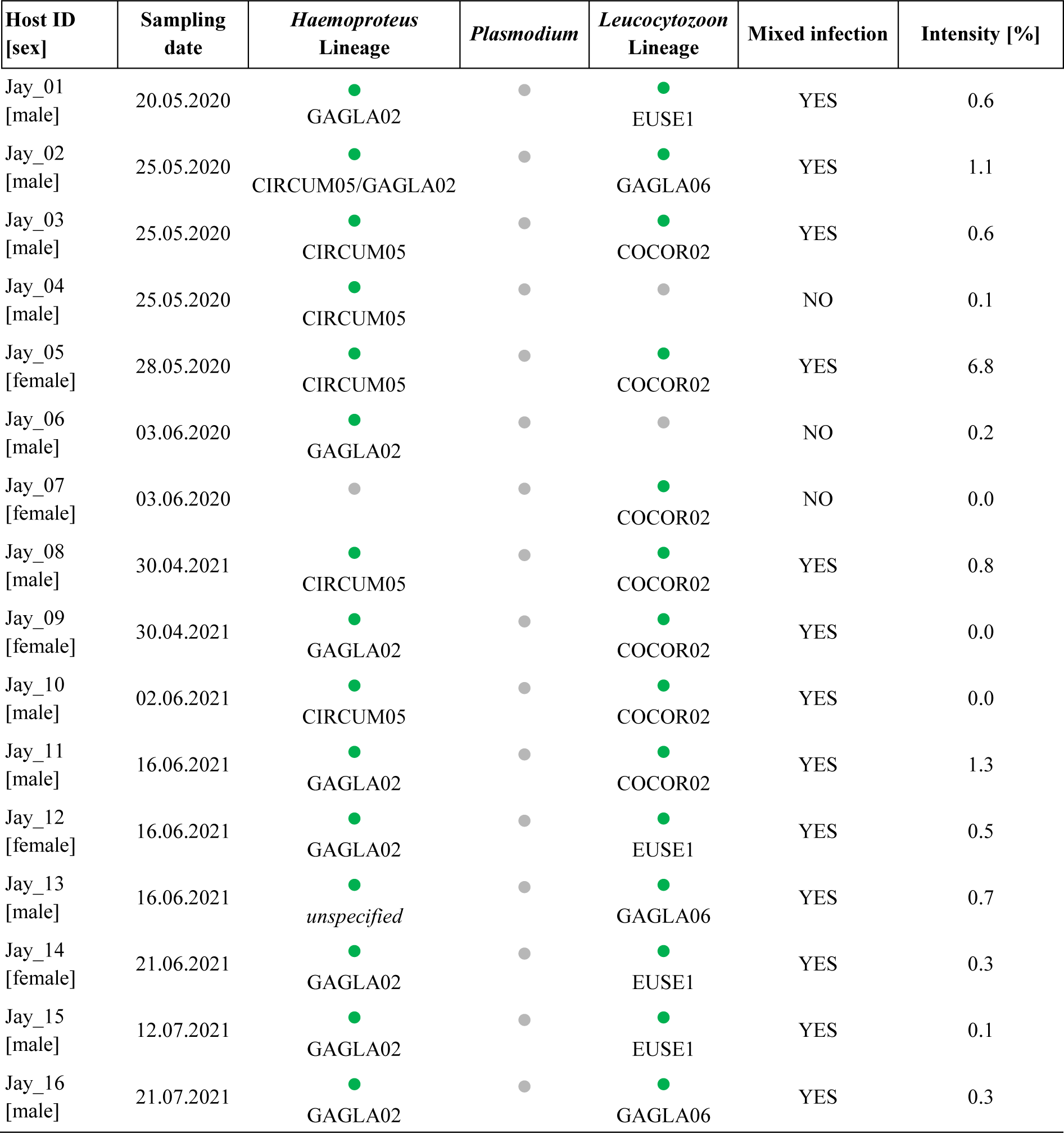
Avian haemosporidian infections in samples Eurasian Jays *Garrulus glandarius*. Infection status refers to nested PCR results (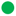 = present infection, 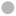 = no detected infection; for infected individuals the respective lineage is given) and infection intensity (percentage of infected erythrocytes) to blood smear counts.

For DNA isolation, we cut a 3 × 3 mm piece of the samples in the FTA card to extract DNA (ammonium-acetate protocol; Martínez et al. 2009). We determined DNA concentration and quality using a NanoDrop2000c UV-Vis spectrophotometer (NanoDrop Technologies, USA).

### Parasite detection

We determined the presence or absence of avian haemosporidians through a nested PCR protocol targeting a 479-base pair (bp) region of the cytochrome *b* gene (cyt *b*; for detailed PCR protocol see Hellgren et al. 2004). In each PCR run, we included DNA from birds with known haemosporidian infection and deionized water as positive and negative controls, respectively. We rerun each sample resulting in a negative PCR reaction to confirm the parasite absence. We visualized the PCR amplicons using QIAxcel Advanced (QIAGEN) high-resolution capillary gel electrophoresis. Subsequently, we sent all PCR products of samples rendering a clear band during gel electrophoresis for Sanger bidirectional sequencing at Microsynth-Seqlab (Sequence Laboratories Goettingen GmbH, Germany). Using CLC Main Workbench 7.6.4 (CLC Bio, Qiagen, Denmark), we assembled and trimmed forward and reverse sequences. To identify lineages, we aligned the sequences with sequences deposited in the MalAvi database using BLASTN 2.3.0+ (Bensch et al. 2009). If the sequencing quality did not allow a clear determination, we repeated the PCR and sequencing. However, regardless this, we could not clearly determine the lineage of one *Haemoproteus*-positive sample (Table 1).

We constructed a haplotype network of haemosporidian lineages (Fig. 1), using the medium joining network method implemented in PopART 1.7 (Leigh & Bryant 2015). The network covers all sequences clearly assigned to a lineage from this study (n = 29) and sequences found in Jays deposited in MalAvi (n = 31). However, we had to exclude two sequences, corresponding to the lineages GAGLA01 and GAGLA04, from MalAvi due to being short (< 476 bp) and containing undetermined nucleotides.

**Figure 1.**
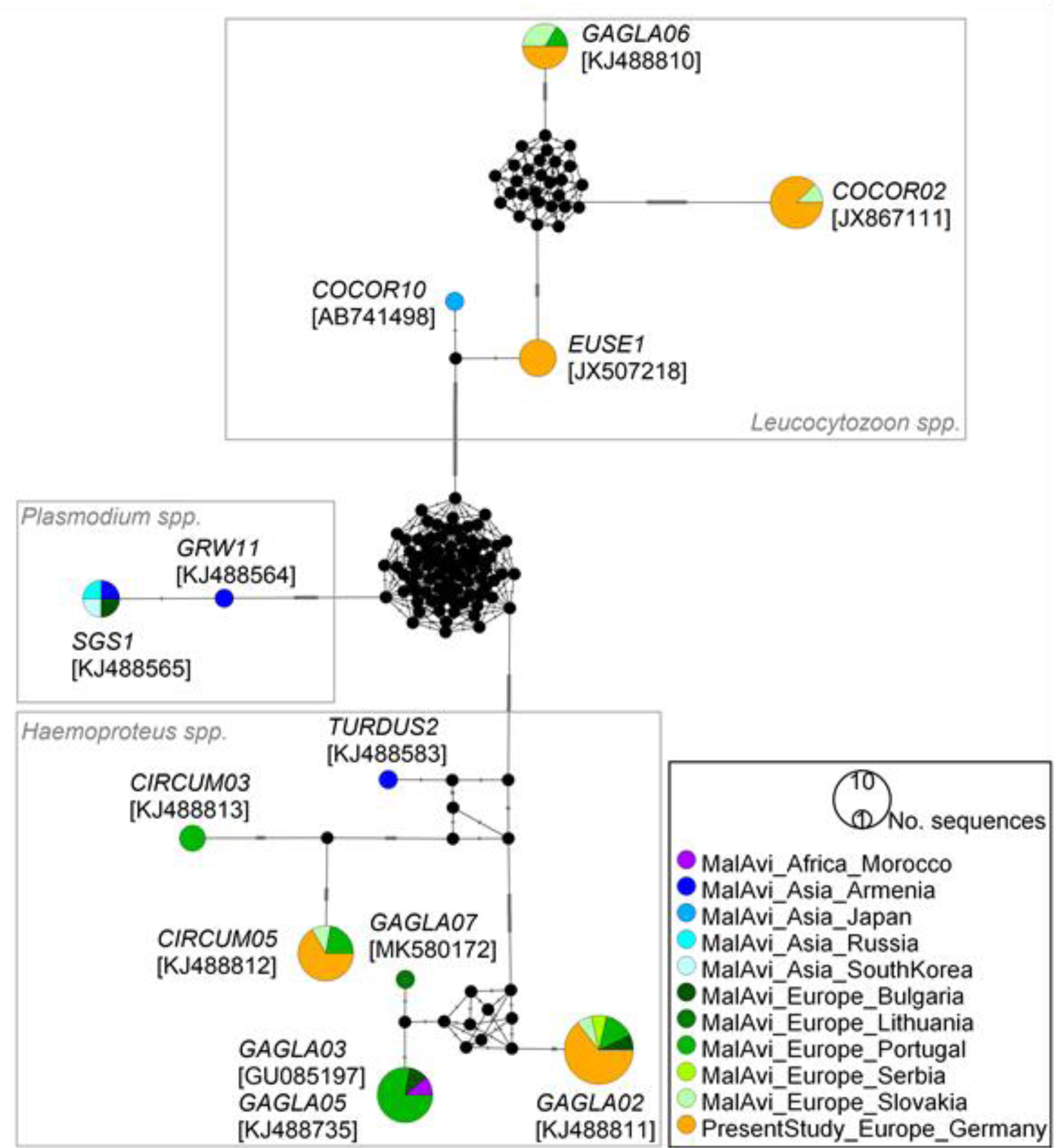
Median-joining network of mitochondrial cytochrome *b* lineages (476 bp, n = 60 sequences) of haemosporidian parasites found in Eurasian Jays *Garrulus glandarius*. Circle size is proportional to the lineage frequency. Lineage names are noted at the associated circles together with one exemplary GenBank association number in parentheses. One hatch mark represents one mutation. Sample origins are represented by different colours, “MalAvi” referring to sequences from other studies deposited in the MalAvi database (MalAvi 2023).

We examined the stained blood smears at × 1000 magnification (light microscope PrimoStar Zeiss, Germany) for at least 10,000 monolayered erythrocytes to calculate infection intensity based on the presence and number of intraerythrocytic gametocytes (e.g. Stanković et al. 2019).

## Results & Discussion

The overall infection rate in the Eurasian Jay population was 100%, i.e. each individual showed a positive PCR result for at least one haemosporidian genus. Interestingly, we could detect no *Plasmodium* infections, while *Haemoproteus* and *Leucocytozoon* infections reached a high prevalence (94% and 88%, respectively), and often occurred as mixed infection (Table 1). However, *Plasmodium* infections (lineages GRW11 and SGS1) have been found in a few Eurasian Jays from Asia and Europe (Beadell et al. 2006; Dimitrov et al. 2010; Drovetski et al. 2014; Fig. 1). Whereas other studies also found no *Plasmodium* infections in Eurasian Jays (Stanković et al. 2019; Šujanová et al. 2021).

Matching our result, a community analysis, including 29 avian species, from the same study area in Hesse found low *Plasmodium* infection prevalence (8%), compared to a higher *Haemoproteus* (68%) and *Leucocytozoon* (60%) prevalence (Strehmann et al. 2023). However, how exactly phylogenetic, climatic, environmental and behavioural factors influence and form the occurrence and distribution of haemosporidian parasites is still under research (e.g. Pigeault et al. 2022; de Angeli Dutra et al. 2023).

The intensity of infection (parasitaemia) varied between 0 and 6.8%, however, most individuals (81%) had an intensity < 1.0% (Table 1). Generally, low-intensity infections (connected with mainly chronic infections) can persist in the hosts without direct visible effects. This fits the observation that none of the sampled Eurasian Jays showed any clinical signs of disease. Yet, some studies showed that also a low-intensity haemosporidian infection can have negative long-term effects on the host fitness, e.g. by influencing reproduction and survival (e.g. Asghar et al. 2015; Schoenle et al. 2017).

We were able to assign the detected *Haemoproteus* infections to two lineages: GAGLA02 (prevalence: 56%, GenBank accession number: OR069477) and CIRCUM05 (38%, OR069478), whereby one Jay individual was infected with both lineages (Table 1). Both lineages already have been proved in Eurasian Jays from other sampling locations in Europe (Fig. 1). Considering the lineage network, the GAGLA-*Haemoproteus* lineages cluster together (Fig. 1). The clustering lineages GAGLA02, GAGLA03 and GAGLA07 have so far only been detected in Eurasian Jays, the lineage GAGLA05 also in another member of the Corvidae family, the Common Raven *Corvus corax* (MalAvi 2023). This suggests a certain host-specificity of these found *Haemoproteus* lineages, which is line with other findings proposing *Haemoproteus* to be rather host-specific (e.g. Ellis et al. 2020; Strehmann et al. 2023). The *Leucocytozoon* lineages GAGLA06 (OR069481) and COCOR02 (OR069479) were so far found in Eurasian Jays and Common Ravens only, too, whereas the *Leucocytozoon* lineage EUSE1 (OR069480) up to now was detected in Common Ravens only (MalAvi 2023). We found four Eurasian Jay individuals to be infected with EUSE1 (Table 1), constituting a new host-lineage interaction. The results show that, similar to the *Haemoproteus* lineages, some *Leucocytozoon* lineages may be quite host-specific to Corvidae species. Therefore, this avian family, which has been rather poorly studied with respect to haemosporidian infections, might be a good model group for further research on haemosporidian host specificity.

## Acknowledgements

We thank Sabine Wagner, Anna Wahle and Wiebke Schäfer for supporting the laboratory work, and Kim Lindner and Sascha Rösner for assistance to the fieldwork.

## Author contributions

YRS, PQ and JFM conceptualized the study and carried out fieldwork. PQ acquired the funding. YRS and NLP performed laboratory work. YRS analysed the data. The first draft of the manuscript was written by YRS. All authors contributed to manuscript revision and approved the submitted version.

## Data availability statement

Sequences are deposited in GenBank (accession numbers OR069477 to OR069481) and will be submitted to the MalAvi database. The (raw) data of this article is freely available for download from PANGAEA (Data Publisher for Earth & Environmental Science) at https://doi.org/10.1594/PANGAEA.961154, https://doi.org/10.1594/PANGAEA.961255, and https://doi.org/10.1594/PANGAEA.961493

## Ethics statement

All applicable institutional and/or national guidelines for the care and use of animals were followed. Animal handling, including an ethical approval according to the national animal protection laws, was carried out under permits of the Regierungspräsidium Gießen, Hesse, Germany (permit number G10/2019).

